# fastISM: Performant *in-silico* saturation mutagenesis for convolutional neural networks

**DOI:** 10.1101/2020.10.13.337147

**Authors:** Surag Nair, Avanti Shrikumar, Anshul Kundaje

## Abstract

Deep learning models such as convolutional neural networks are able to accurately map biological sequences to associated functional readouts and properties by learning predictive *de novo* representations. *In-silico* saturation mutagenesis (ISM) is a popular feature attribution technique for inferring contributions of all characters in an input sequence to the model’s predicted output. The main drawback of ISM is its runtime, as it involves multiple forward propagations of all possible mutations of each character in the input sequence through the trained model to predict the effects on the output. We present fastISM, an algorithm that speeds up ISM by a factor of over 10x for commonly used convolutional neural network architectures. fastISM is based on the observations that the majority of computation in ISM is spent in convolutional layers, and a single mutation only disrupts a limited region of intermediate layers, rendering most computation redundant. fastISM reduces the gap between backpropagation-based feature attribution methods and ISM. It far surpasses the runtime of backpropagation-based methods on multi-output architectures, making it feasible to run ISM on a large number of sequences. An easy-to-use Keras/TensorFlow 2 implementation of fastISM is available at https://github.com/kundajelab/fastISM, and a hands-on tutorial at https://colab.research.google.com/github/kundajelab/fastISM/blob/master/notebooks/colab/DeepSEA.ipynb.

## Introduction

High-throughput experimental platforms have revolutionized the ability to profile diverse biochemical and functional properties of biological sequences such as DNA, RNA and proteins. These datasets have powered highly performant deep learning models of biological sequences that have achieved state-of-the art results for predicting protein-DNA binding, protein-RNA binding, chromatin state, splicing, gene expression, long-range chromatin contacts, protein structure and functional impact of genetic variation (Zhou *et al.*, 2018; Zhou and Troyanskaya, 2015; Jaganathan *et al.*, 2019; Alipanahi *et al.*, 2015; Eraslan *et al.*, 2019; Torrisi *et al.*, 2020; Kelley *et al.*, 2016; Avsec *et al.*, 2019; Kelley *et al.*, 2018; Koo *et al.*, 2018; Fudenberg *et al.*, 2020).

Convolutional neural networks (CNNs) are widely used for modeling regulatory DNA since they are well suited to capture known properties and invariances encoded in these sequences (Kelley *et al.*, 2016; Alipanahi *et al.*, 2015; Zhou and Troyanskaya, 2015). CNNs map raw sequence inputs to binary or continuous outputs by learning hierarchical layers of *de-novo* motif-like pattern detectors called convolutional filters coupled with non-linear activation functions. Recurrent neural networks (RNNs) (Hochreiter and Schmidhuber, 1997) are another class of sequential models that have been very effective for modeling protein sequences (Torrisi *et al.*, 2020). However, RNNs and hybrid CNN-RNN architectures have only shown moderate performance improvements for modeling regulatory DNA (Quang and Xie, 2016; Hassanzadeh and Wang, 2016; Shen *et al.*, 2018). Compared to recurrent architectures, CNNs also have the advantage of being more computationally efficient and easily interpretable. For example, convolutional filters are reminiscent of classical DNA motif representations known as position weight matrices (PWMs) (Trabelsi *et al.*, 2019). Hence, CNNs continue to be the most popular class of architectures for modeling regulatory DNA sequences (Eraslan *et al.*, 2019).

A primary use case for these deep learning models of regulatory DNA is to decipher the *de novo* predictive sequence features and higher-order syntax learned by the models that might reveal novel insights into the regulatory code of the genome. Hence, several feature attribution methods have been developed and used to infer contribution scores (or importance scores) of individual characters in input sequences with respect to output predictions of neural network models such as CNNs. A popular class of feature attribution methods use backpropagation to efficiently decompose the output prediction of a model, given an input sequence, into character-level attribution scores (Shrikumar *et al.*, 2017; Sundararajan *et al.*, 2017; Lundberg and Lee, 2017; Simonyan *et al.*, 2013). The gradient of the output with respect to each observed input character — commonly referred to as a saliency map (Simonyan et al., 2014)— is one such method for attributing feature importance. Other related approaches such as DeepLIFT (Shrikumar *et al.*, 2017) and integrated gradients (Sundararajan *et al.*, 2017; Jha *et al.*, 2020) modify the backpropagated signal to account for saturation effects and improve sensitivity and specificity. These attribution scores can be used to infer predictive subsequences within individual input sequences which can then be aggregated over multiple sequences to learn recurring predictive features such as DNA motifs (Shrikumar *et al.*, 2018).

*In-silico* Saturation Mutagenesis (ISM) is an alternate feature attribution approach that involves making systematic mutations to each character in an input sequence and computing the change in the model’s output due to each mutation. ISM is the computational analog of saturation mutagenesis experiments (Patwardhan *et al.*, 2009) that are commonly used to estimate the functional importance of each character in a sequence of interest based on its effect size of mutations at each position on some functional read out. ISM is the de-facto approach to predict the effects of genetic variants in DNA sequences (Zhou and Troyanskaya, 2015; Kelley *et al.*, 2016; Wesolowska-Andersen *et al.*, 2020).

In the context of computing feature attributions with respect to a single scalar output of a model, ISM can be orders-of-magnitude more computationally expensive than backpropagation-based feature attribution methods, since it involves a forward propagation pass of the model for every mutation of every position in an input sequence (Eraslan *et al.*, 2019). By contrast, backpropagation-based methods can compute attribution scores of all possible characters at all positions in an input sequence in one or a few backward propagations of the model. The inefficiency of ISM is particularly onerous when ISM is performed on a large number of sequences or for a large number of models. For example, to obtain robust variant effect prediction, (Wesolowska-Andersen *et al.*, 2020) recently trained 1000 convolutional neural networks to predict variants in chromatin regulatory features of pancreatic islets, and averaged the ISM scores over the trained models to confer robustness against heterogeneity that stems from different random parameter instantiations of the same model at the beginning of the training process (Wesolowska-Andersen *et al.*, 2020).

However, despite this gap in efficiency, ISM does offer some salient benefits over backpropagation-based methods. In comparison to most backpropagation-based methods that often use heuristic rules and approximations, ISM faithfully represents the model’s response to mutations at individual positions. This makes it the method of choice when evaluating the effect of genetic variants on the output (Zhou and Troyanskaya, 2015; Zhou *et al.*, 2018; Wesolowska-Andersen *et al.*, 2020), and it is also used as a benchmark reference when evaluating fidelity of other feature attribution methods (Koo and Ploenzke, 2020). Unlike ISM, backpropagation-based methods like DeepLIFT and Integrated Gradients rely on a predefined set of “neutral” input sequences that are used as explicit references to estimate attribution scores. The choice of reference sequences can influence the scores and so far the selection of reference sequences has been ad-hoc (Jha *et al.*, 2020; Eraslan *et al.*, 2019). ISM also has some benefits for models with a large number of scalar or vector outputs since each forward propagation performed during ISM reveals the impact of a single mutation on every output of the model. For example, massively multi-task models are quite popular for mapping regulatory DNA sequences to multiple molecular reads outs in large collections of biosamples (Zhou *et al.*, 2018; Zhou and Troyanskaya, 2015; Jaganathan *et al.*, 2019; Alipanahi *et al.*, 2015; Eraslan *et al.*, 2019). Further, a recent class of models called profile models have been developed to map regulatory DNA sequences to vector outputs corresponding to quantitative regulatory profiles (Avsec *et al.*, 2019; Kelley *et al.*, 2018). These models output a vector of signal values often at base-resolution that can be as long as the input sequence. ISM can reveal how perturbing individual nucleotides in the input alters the signal across all positions in the output profile. By contrast, backpropagation-based importance scoring methods would need to perform a separate backpropagation for every output position in order to estimate comparable feature attributions, which would linearly increase the computational cost in the number of outputs. For these reasons, a computationally efficient implementation of ISM would be attractive.

We introduce fastISM, an algorithm that speeds up ISM for CNNs. fastISM is based on the observation that CNNs spend the majority of computation at prediction time in convolutional layers and that single point mutations in the input sequence affect a limited range of positions in intermediate convolutional layers. fastISM restricts the computation in intermediate layers to those positions that are affected by the mutation in the input sequence. fastISM cuts down the time spent in redundant computations in convolution layers at positions that are unaffected by mutations in the input, resulting in significant speedups.

We provide a fully functional and well-tested package implementing the fastISM algorithm for Keras models in TensorFlow 2. We benchmark the speedup obtained by running fastISM on a variety of architectures and show that fastISM can achieve order-of-magnitude improvements over standard ISM implementations. fastISM reduces the gap between ISM and backpropagation-based methods in terms of runtime on single-output architectures, and far surpasses them on multi-output architectures.

## Results

### *In-silico* saturation mutagenesis for CNNs is bottlenecked by redundant computations in convolution layers

We motivate fastISM by using an example based on the multi-task Basset model (Kelley *et al.*, 2016) that maps DNA sequences to binary labels of chromatin accessibility across 100s of cell types and tissues (tasks). The Basset model consists of 3 convolution layers of kernel sizes 19, 11 and 7, which are followed by max pool layers of size 3, 4 and 4 respectively. The output after the 3rd convolution and max pool is flattened. This is followed by 2 fully connected layers with 1000 hidden units, and a final fully connected layer that predicts the outputs. We consider a slight modification of the original architecture that operates on 1000bp input sequence and has 10 binary scalar outputs. In addition, all convolution layers are assumed to be padded such that the length of the output sequence is the same as the input to the layer (**Fig 1**).

**Fig 1.**
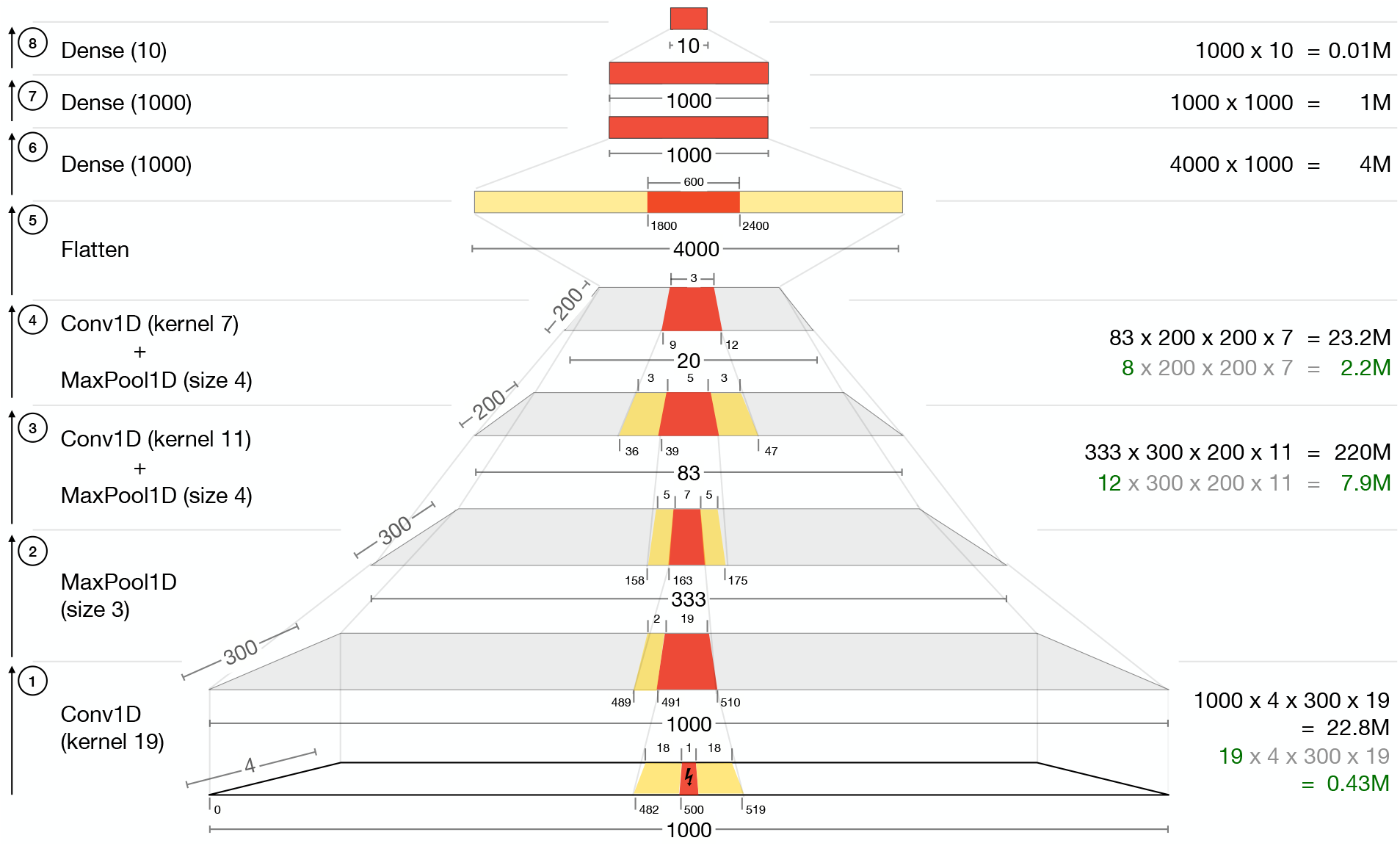
Annotated diagram of a Basset-like architecture (Kelley *et al.*, 2016) on an input DNA sequence of length 1000, with a 1 base-pair mutation at position 500 (0-indexed). Positions marked in red indicate the regions that are affected by the point mutation in the input. Positions marked in yellow, flanking the positions in red, indicate unaffected regions that contribute to the output of the next layer. Ticks at the bottom of each layer correspond to position indices. Numbers on the right in black indicate the approximate number of computations required at that layer for a standard implementation of ISM. For convolution layers, the numbers in gray and green indicate the minimal computations required. We omit layers such as activations and batch normalization for simplicity, as they do not change the affected regions.

For a single forward propagation through the network, the majority of compute time is spent in the convolutional layers. For a given convolution layer, the number of computations is proportional to kernel size, input filters, output filters and output length. For a fully connected layer, it is proportional to input and output lengths. The numbers in black in **Fig 1** show the approximate computations required at each convolution and fully connected layer for a single forward propagation. We ignore the computation spent in intermediate layers such as activations, batch normalization and max pool, since they are dominated by the computational cost of convolutional and fully connected layers. The 3 convolutional layers require approximately 23M, 220M and 23M computations respectively, while the 3 fully connected layers require approximately 4M, 1M and 0.01M computations respectively. Thus, the computations required in the convolutional layers combined exceed that of the fully connected layers by a factor of 50x. In practice, we timed the convolutional layers and fully connected layers and observed that the factor is closer to 40x for the same architecture.

For a given reference input sequence, a simple implementation of ISM involves highly redundant computations. Typically, ISM is implemented by inserting mutations at each position in the input one at a time and making a forward pass through the entire model using the perturbed sequences as inputs. However, local perturbations in the input only affect local regions in intermediate convolutional layers, while regions farther away remain identical to their values for the reference (unperturbed) input sequence.

For each layer, the regions that are affected by the single base pair mutation in the input, and the minimal regions required to compute the output of the next layer are shown in **Fig 1**. Consider a single base pair mutation in the middle of the 1000 bp input sequence at position 500 (0-indexed). The first convolution (layer 1) has a kernel size of 19 and the input sequence is padded with zeros at 9 positions on both sides. The output sequence length is thus 1000. As the convolution filter scans across the input sequence, the mutation at position 500 will be involved in 19 contiguous output positions of the next layer from positions 491-510. None of the other 1000-19 outputs will be affected by the mutation, and computing them is redundant. 18 positional inputs on either side of the mutation will be involved in generating the 19 contiguous outputs of the next layer. The first max pool (layer 2) has a size of 3, and the output length is 333. The affected input region from 491-510 will only affect ⌊491/3⌋-⌊510/3⌋, i.e. 163-170 in the output of the max pool region. However, in order to compute the 163rd output, the max pool would also require the convolution output values at positions 489 and 490, which would be the same as the values for the reference (unperturbed) input sequence.

The above exercise can be extended to the next two convolution layers. For simplicity, the convolution and max pool layers are combined. The output of the first max pool affects positions 163-170. The next convolution has a kernel size of 11 and a padding of zeros at 5 positions on both sides. The output of the convolution followed by max pool (layer 3) affects 5 out of 83 positions, and the output of layer 4 affects 3 out of 20 positions. This output is then reshaped to a single vector, which is then fed through 3 fully connected layers. By design, a single mutation in the input sequence has the potential to affect every position in the fully-connected layer; thus, the activations of all subsequent layers in the network must be recomputed for the mutated input, as is the case with standard ISM.

Given that the majority of computation occurs before the fully-connected layer, the actual computations required to track the effect of a single base-pair mutation in the input are much smaller than the total computations in a standard forward propagation. The values on the right in green and gray in **Fig 1** show the minimum number of computations required such that computations are only restricted to the regions affected by the mutation in the input. For saturation mutagenesis in which the above operations are repeated for perturbations at all positions, the amount of redundant calculations adds up and contributes to ISM’s unfavourable runtime.

Since the majority of activations prior to the fully-connected layers remain unchanged by a single-base mutation, the activations of these layers on the unperturbed sequence can be computed once and reused when running ISM over the different positions in the input sequence. By restricting ISM computations to only positions affected by the input mutation at each layer, the number of computations can theoretically be reduced from approximately 23 + 220 + 23 + 4 + 1 M = 271M to 0.5 + 8 + 2 + 4 + 1 M = 15.5M computations, down by a factor of 17. In practice, there may be overheads from other steps such as concatenating the unperturbed flanking regions at each intermediate layer, that would dampen the realised speedup.

These observations suggest that it should be possible to define a custom model that performs only the required computations for a mutation at each input position. However, it would be cumbersome to write an architecture specifically for the purpose of ISM for each model, and compute the positions required at each intermediate layer for a specific input mutation. Hence, we developed fastISM, a method to speed up ISM by leveraging the above mentioned redundancies without requiring any explicit re-specification of the model architecture by the user (Supplementary Methods).

### fastISM yields order-of-magnitude improvements in speed for different architectures

fastISM takes as input any sequence model and first reduces it to a computational graph representation. It chunks the graph into appropriate segments that can each be run as a unit. The model is augmented to return intermediate outputs at the end of each segment for unperturbed input sequences. For a given set of input sequences, these intermediate outputs are cached. A second model is then initialized, that largely resembles the original model, but incorporates mechanisms to concatenate slices of the cached unperturbed intermediate outputs with the affected intermediate outputs, and compute each layer’s output on the least required input. fastISM processes a group of input sequences at a time. For each group of sequences, fastISM is run on multiple batches such that sequences in a given batch are all perturbed at the same position.

We benchmarked fastISM against a standard implementation of ISM. We choose 3 types of models that take DNA sequence as input— the Basset architecture (Kelley *et al.*, 2016), the Factorized Basset architecture (Wnuk *et al.*, 2019) and the BPNet architecture (Avsec *et al.*, 2019). The first two models output scalar values for each output task, whereas the BPNet model outputs a profile vector of length equal to the input sequence length, and a scalar count. ISM is performed by recording the outputs for all 3 alternate mutations at each position. We benchmark the three models for 1000bp and 2000bp length inputs.

We also compare fastISM to three backpropagation-based feature attribution methods — Gradient x Input (input masked gradient), Integrated Gradients (Sundararajan *et al.*, 2017), and DeepSHAP (Lundberg and Lee, 2017). DeepSHAP is an extension of the DeepLIFT algorithm that is implemented by overriding tensorflow gradient operators; we used DeepSHAP because it has a more flexible implementation than the original DeepLIFT repository. We set the number of steps for Integrated Gradients to 50 and a single default reference of all zeros, and the number of dinucleotide reference sequences for DeepSHAP to 10. For models with a single scalar output, the backpropagation-based methods are run with respect to the scalar output. For BPNet, the methods are run with respect to each output in the profile vector as well as the scalar output. While ISM returns one value (change in output score) for each of the 3 alternative nucleotides (with respect to the observed nucleotide) at each position; Integrated Gradients, DeepSHAP and Gradients return one value for each of the 4 possible nucleotides at each position.

The results are summarised in **Table 1**. For the Basset and Factorized Basset architectures, fastISM speeds up ISM by more than 10x when computing importance scores for a single output task, and the speedup increases with increasing input sequence length. This is expected since, for a fixed architecture, the length of the affected regions in the convolutional layers are independent of input sequence length. Remarkably, fastISM runtimes, though slower than Gradient x Input, are competitive with runtimes of Integrated Gradients (within 2x) and DeepSHAP (within 4x) for single scalar output models. Also note that fastISM and ISM provide importance scores with respect to every output task; computing scores for multiple output tasks while the backpropagation-based methods would multiply their runtime by the number of output tasks.

**Table 1:** Comparison of fastISM with standard ISM, Gradient x Input, Integrated Gradients with 50 steps and a single all-zeros reference, and DeepSHAP with 10 references for 3 different models with 1000bp and 2000bp length inputs. All times in seconds per 100 input sequences. Time relative to fastISM in parentheses. For BPNet models which output a profile vector as well as a count scalar, Gradient x Input, Integrated Gradients and DeepSHAP were computed in a loop with respect to each output of the profile and the count scalar (*).

| Architecture | Layers | Input Size | Outputs | fastISM | Standard ISM | Gradient x Input | Integrated Gradients | DeepSHAP |
| --- | --- | --- | --- | --- | --- | --- | --- | --- |
| Basset | 3 Conv + 3 max pool, 3 fully connected | 1000 | Single scalar | 2.70 | 27.36 (10.1) | 0.04 (<<1) | 2.34 (0.8) | 1.75 (0.7) |
|  |  | 2000 |  | 6.49 | 100.44 (15.4) | 0.08 (<<1) | 4.61 (0.7) | 3.03 (0.4) |
| Factorized Basset | 9 Conv + 3 max pool, 3 fully connected | 1000 | Single scalar | 5.47 | 68.97 (12.6) | 0.09 (<<1) | 4.82 (0.9) | 2.64 (0.5) |
|  |  | 2000 |  | 18.04 | 262.24 (14.5) | 0.17 (<<1) | 9.47 (0.5) | 4.63 (0.25) |
| BPNet | 2 Conv, 9 Dilated Conv, skip connections | 1000 | Profile (length 1000 vector) + scalar | 28.97 | 46.09 (1.6) | 41.49* (1.4) | 4399* (151) | 1743* (60) |
|  |  | 2000 | Profile (length 2000 vector) + scalar | 81.52 | 173.96 (2.1) | 126.41* (1.5) | 12440* (152) | 6427* (78) |

The speedup of fastISM for the BPNet architecture relative to standard ISM is more modest— 1.6x for 1000bp input and 2.1x for 2000bp input. This can be attributed to dilated convolutions; since the BPNet architecture includes dilated convolutions with an exponentially increasing dilation rate, the receptive fields for the later dilated convolutions are very large. As a result, the regions affected by a single base pair change in the input span a sizable fraction of intermediate layers, and the computations involved beyond those layers approach those of a standard implementation.

For the BPNet architecture which outputs a profile vector, if one is interested in attributing the predicted value at each position in the output vector to the input nucleotides, one would need to run the backpropagation methods for every output position, which drastically slows them down. DeepSHAP and Integrated Gradients take over 50x time of the fastISM implementation. Thus, fastISM speeds up ISM by an order-of-magnitude and narrows the gap in compute time between backpropagation-based methods and ISM.

## Conclusions

*In-silico* saturation mutagenesis (ISM) is an important post-hoc feature attribution method that has gained applicability as a tool to interpret deep learning models for genomics and to interrogate the effect of variants. ISM has largely been treated as a static method with unfavourable time complexity compared to more recent backpropagation-based model interpretability methods. We challenge this notion by introducing fastISM, a performant implementation of ISM for convolutional neural networks. fastISM leverages the simple observation that the majority of computations performed in a traditional implementation of ISM are redundant. fastISM improves runtime of ISM by over 10x for commonly used convolutional neural networks, and a factor of 2x for profile networks with exponentially wide dilated convolutions. This brings down ISM’s runtime in the ballpark of backpropagation-based methods such as Integrated Gradients and DeepSHAP for single-output models, and dramatically surpasses the runtime of backpropagation-based methods for multi-output methods, making it more feasible to run ISM genome-wide and on a large number of models.

## Notes

https://github.com/kundajelab/fastISM

https://colab.research.google.com/github/kundajelab/fastISM/blob/master/notebooks/colab/DeepSEA.ipynb

### Competing Interest Statement

The authors have declared no competing interest.

