## Supplementary Methods for "fastISM: Performant *in-silico* saturation mutagenesis for convolutional neural networks"

**fastISM Algorithm Details** **2**

Obtaining a Simplified Computational Graph from the Keras Model 3

The Slice-Assign Operation 4

Segmenting the Computational Graph 4

Constructing the Intermediate Output (IntOut) Model 5

Perturbed Range Calculations for Segments 6

Constructing the Mutation Propagation (MutProp) Model 6

Execution on Input Sequences 7

Handling Additional Non-sequence Inputs 7

**Benchmarking** **8**

**fastISM Implementation** **10**

[1]Department of Computer Science, Stanford University [2]Department of Genetics, Stanford University

**fastISM Algorithm Details**

fastISM builds upon the observation that local perturbations in the input sequence tend to affect only local regions in the convolutional layers. These regions are narrower in the earlier convolutional layers, and grow wider with increasing depth. Given a mutation at a fixed position in the input sequence, standard implementations recompute intermediate outputs from unperturbed regions farther away from the mutation that do not change after introducing the mutation. fastISM computes these intermediate outputs once for each reference unperturbed input sequence and caches them. For each different positional mutation in the input, appropriate windows around perturbed regions can be reused at each intermediate layer. fastISM restricts computation in convolutional layers to only the affected regions and unperturbed flanking regions around it, which typically depend on kernel width and dilation rate.

fastISM takes a Keras model as input. The main steps of fastISM are as follows:

- One-time Initialization:
    - Obtain the computational graph from the model and chunk it into segments that can be run as a unit
    - Augment the model to create an “intermediate output model” (IntOut model) that returns intermediate outputs at the end of each segment for reference input sequences

[1]Department of Computer Science, Stanford University [2]Department of Genetics, Stanford University

    - Create a second “mutation propagation model” (MutProp model) that largely resembles the original model, but incorporates as additional inputs the necessary flanking regions from outputs of the IntOut model on reference input sequences between segments
- For each batch of input sequences:
    - Run the IntOut model on the sequences (unperturbed) and cache the intermediate outputs at the end of each segment
    - For each positional mutation:
        - Introduce the mutation in the input sequences
        - Run the MutProp model feeding as input appropriate slices of the IntOut model outputs

Each of these steps is described in more detail in the following subsections.

*Obtaining a Simplified Computational Graph from the Keras Model*

Keras allows defining models through their Sequential and Functional API. Models can also be recursively defined and used as building blocks of other models. This necessitates a representation that is easier to manipulate and independent of how the model is specified by the user. The input Keras model is converted to a graph representation. The method runs recursively on sub-models (if any), and finally returns a bipartite graph that consists of tensors and Keras layers. A tensor is derived from a single layer (in-degree of 1). While most layers also take in one input tensor, certain aggregation layers such as Add can have an in-degree greater than 1. Currently, a limited set of Keras layers are supported that span the most commonly used operations for sequence based models.

***The Slice-Assign Operation***

Slice-Assign operation is a custom operation written in Tensorflow that takes in two tensors `x` and `y`, an index `i` and performs the operation `x[:,i:i+y.shape[1]] = y`. Thus it overlays `y` on `x` starting from index `i`. Typically, `x` is a slice of unperturbed intermediate output at a particular layer (from the IntOut model), and `y` is the perturbed tensor at the intermediate layer. The MutProp model that performs the main ISM computation given a perturbed input relies on the Slice-Assign operation to stitch the perturbed regions ( `y` ) with the unperturbed regions ( `x` ).

***Segmenting the Computational Graph***

The computational graph is separated into segments that are run as a unit. The underlying idea is that the perturbed region will be overlaid on the unperturbed intermediate outputs from the IntOut models only at the junction of two segments through a Slice-Assign operation, and not within a segment. Consider for example a convolutional layer followed by an activation layer and then a batch normalization layer. For the activation and batch normalization layers, the output at the $i^{th}$ position depends only on the input at the $i^{th}$ position. Hence, they would not require any additional unperturbed flanking intermediate outputs for their computation. These two layers can thus be merged with the preceding convolution layer into a single segment. This would keep the number of intermediate outputs that are cached to a minimum, as well as the number of Slice-Assign operations required.

In general, max pool layers may require additional unperturbed flanking intermediate outputs. As seen in the first max pool layer in **Fig 1** which has a size of 3, the max pool requires two additional positions at its left edge. This would necessitate that the max pool layer be in a different segment as the preceding convolution. However, using a simple modification, the max pool layer and the preceding convolution can be merged into a single segment. Given an input mutation region, coordinates can be computed such that the output of the convolution will be sufficient for the max pool layer. In the example of **Fig 1**, this would mean that for a mutation at index 500 in the input sequence, instead of providing nucleotides from positions 482-519 as input, the region can be expanded two bases to the left to span 480-519. As a result, the output of the first convolution will span 489-510, and the max pool layer would not require any additional inputs. This adds a slight increase to the computations performed by the convolution layer, but decreases the number of intermediate outputs stored and Slice-Assign operations.

Thus, in general, convolution layers are merged with other “see-through layers” (such as activations and batch normalization) as well as max pool layers into one segment. Every new convolution layer starts a new segment. Aggregation layers that take multiple inputs, such as Add, also start a new segment. Once a “stop layer”, such as Flatten or Reshape layer is reached, it is assumed that all positions of downstream layers are affected by the mutation. Thus, before a stop layer, the Slice-Assign operation overlays the perturbed regions over the entire unperturbed intermediate output at that layer. All downstream computations then are equivalent to that in a standard ISM implementation.

*Constructing the Intermediate Output (IntOut) Model*

Once the graph is segmented, the tensors in the computational graph that lie at the junction of two segments are identified. A new IntOut model is created that resembles the input model, but returns as additional outputs the tensors that are at the junction of two segments. The weights of all layers are set to be equal to those of the input model.

*Perturbed Range Calculations for Segments*

The first segment begins with the input sequence tensor. Typically, a mutation is introduced at each position in the input sequence, and this gives us a list of single-character ranges that are perturbed. Starting with this list of perturbed ranges, we can recursively compute the ranges of the output perturbed regions for all segments, as well as the minimal range of the input required to compute the corresponding output. This minimal range is used to slice the unperturbed intermediate output that is combined through a Slice-Assign operation with the perturbed output of the previous segment. As seen in **Fig 1**, for the first segment that would comprise the first convolution and max pool layers, the perturbed input region is 500-501. Providing input sequence from 480-519 (explained in section on *Segmenting the Computational Graph*), the output perturbed range at the end of the max pool layer would span 163-170.

To achieve this, we work out rules for convolutional and max pool layers. For each of these layers, given an input range of perturbed region, we compute the range of the flanking unperturbed region required, and the corresponding output perturbed region indices, taking special care when the perturbed ranges are close to the beginning or end of the sequence. Given multiple layers within a segment, these values can be chained together to obtain the range of flanking unperturbed region and output perturbed region indices for the segment. For aggregation layers as Add layers that take multiple inputs, the input perturbation region is chosen such that it spans the perturbation regions of all inputs.

*Constructing the Mutation Propagation (MutProp) Model*

The MutProp model largely resembled the original input model. At the junctions of segments, two additional inputs are added to the MutProp model— the slice of unperturbed output from the IntOut model at that segment, and a scalar index starting at which the perturbed output of the previous layer is overlaid on the unperturbed output. These are combined through a Slice-

Assign operation. The weights of all layers are set to be equal to those of the input model. Notably, the padding of all convolutional layers is set to 0 since the models run on subsets of the entire tensor.

***Execution on Input Sequences***

fastISM runs on batches of input sequences, all of which are mutated at all specified positions in the input. For each batch of input sequences, the IntOut model is run once on the sequences (unperturbed) and the intermediate outputs returned from the end of each segment are cached. For each positional mutation, the MutProp model is run by feeding as input slices of the IntOut model's outputs (after appropriate padding) and the indices required for the Slice-Assign operations.

***Handling Additional Non-sequence Inputs***

fastISM also allows models that take in additional inputs apart from the primary sequence input which is mutated. All nodes in the graph that are exclusively derived from secondary non-sequence input are assumed to be constant through the ISM. When the graph is segmented, secondary non-primary sequence inputs are processed after primary sequence input. Descendant nodes of non-sequence inputs that merge with descendants of the sequence input before a "stop layer" (such as Flatten, Reshape and Dense, after which all positions of downstream layers are assumed to be affected by the input mutation) are not allowed in the current implementation. For nodes that merge with sequence input after a stop layer (e.g. through a Concatenation), the output before merging is cached and reused. This is another way of avoiding redundant computation, as these computations that operate on non-sequence input that are unaffected by the mutation will not be repeated for each mutation.

**Benchmarking**

In order to benchmark fastISM, we compare it with a standard implementation of ISM. The standard implementation involves incorporating a mutation in the batch of sequences and running the model on the perturbed sequence, for all mutations. We benchmark the models on 3 architectures:

1. Basset architecture which consists of 3 sets of convolutions + max pool layers, followed by 3 fully connected layers
2. Factorized Basset architecture which is similar to the Basset architecture except the 3 convolutions are replaced by multiple convolutions with narrower kernels. In total, it consists of 9 convolutions, 3 max pool layers and 3 fully connected layers
3. BPNet architecture, which performs an initial convolution on the input sequence, followed by 9 dilated convolutions for which the dilation rate increases as $2^i$ where i corresponds to the $i^{th}$ dilation. Skip connections are added such that input of each dilation layer is added to its output. The model has two outputs: the first is a transposed convolution on the flattened output of the dilated convolutions, the second is a fully connected layer on top of Global Average Pool output of the dilated convolutions. The transposed convolution outputs a profile vector, while the fully connected layer outputs a count scalar.

We notice that fastISM tends to exhibit overhead costs and is slow for small batch sizes. Hence we swept batch sizes from as small as 64 to as large as 4096 that could fit in GPU memory for both fastISM and the standard implementations. For a batch of input sequences, we introduced all 3 alternate mutations at the $i^{th}$ position. We recorded the best times per 100 examples across these batch sizes. All benchmarking is performed on random input tensors with randomly initialized models.

We also ran three backpropagation-based methods for comparison for the same models. We used a version a few commits after v0.36.0 of DeepSHAP (https://github.com/slundberg/shap/tree/f603b3287), v0.5.4 of the library Alibi for Integrated Gradients (https://github.com/SeldonIO/alibi/tree/v0.5.4) and a simple custom implementation of Gradient x Input. Input batch sizes for DeepSHAP and Gradient x Input, and internal batch sizes for Integrated Gradients were varied and the best time per 100 examples was recorded. We used 50 steps for Integrated Gradients (with a single all-zeros reference; runtime would linearly increase if more references were used) and 10 dinucleotide shuffled sequences as reference DeepSHAP. Note that for these methods, importance scores are computed for each character at each position in the input. fastISM and standard ISM implementations introduce 3 alternate mutations at each position and thus return three values for each position in the input sequence.

Regarding the runtime of DeepSHAP: in the DeepSHAP implementation, each batch consists of a single input sequence and its corresponding reference values; thus, batch sizes are not as ideal as in the case where the importance scores for multiple input sequences could be calculated in the same batch. Further, the DeepSHAP implementation duplicates the input sequences to align with the number of references (https://github.com/slundberg/shap/blob/f603b3287/shap/explainers/_deep/deep_tf.py#L298); these aspects contribute to the slower runtime of DeepSHAP relative to a hypothetical ideal implementation. We used DeepSHAP because the implementation can work with diverse model architectures and supports dynamic references.

All experiments were performed on a CentOS Linux release 7.6.1810 machine with 128GB RAM. A single NVIDIA Tesla P100-PCIE-16GB GPU was used, with CUDA v10.1.168, CUDNN v7.6.4, and TensorFlow v2.3.0. fastISM v0.4.0 (https://github.com/kundajelab/fastISM/tree/v0.4.0) was used for all benchmarks.

**fastISM Implementation**

fastISM has been implemented in TensorFlow v2.3.0 and supports Keras sequence models. The implementation is available at https://github.com/kundajelab/fastISM. fastISM provides a simple to use API for performing ISM:

```python
from fastism import FastISM

# any Keras model
fast_ism_model = FastISM(model)

for seq_batch in sequences:
    # seq_batch has dimensions (batch_size, seq_length, alphabet_size)
    ism_seq_batch = fast_ism_model(seq_batch)
    # ism_seq_batch has dimensions (batch_size, seq_length, num_outputs)
```

When fastISM is initialized on an input model, it checks by default if the output for a small batch of sequences matches that of a simple non-optimized implementation of ISM. This ensures that any internal inconsistencies in the implementation are caught before the user runs fastISM on multiple sequences.

It also provides support for multiple-input models, choosing different ranges to perturb in the input and specifying the perturbation values. Full functionality is documented in the repo.
